# Migrastatic Therapy as a Potential Game-Changer in Adaptive Cancer Treatment

**DOI:** 10.1101/2024.12.28.630610

**Authors:** Katharina Schneider, Louise Spekking, Sepinoud Azimi, Barbora Peltanová, Daniel Rösel, Joel S. Brown, Robert A. Gatenby, Jan Brábek, Kateřina Staňková

**Affiliations:** Maastricht University; Delft University of Technology; Charles University; Charles University, Faculty of Science; Moffitt Cancer Center; Faculty of Science, Charles University

**Keywords:** Migrastatics, adaptive therapy, spatial modeling, game theory

## Abstract

Adaptive therapy, which anticipates and counters the evolution of resistance in cancer cells, has gained significant traction, especially following the success of the Zhang et al.’s protocol in treating metastatic castrate-resistant prostate cancer. While several adaptive therapies have now advanced to clinical trials, none currently incorporates migrastatics, i.e. treatments designed to inhibit cancer cell metastasis.

In this study, we propose integrating migrastatics into adaptive therapy protocols and evaluate the potential benefits of using a game-theoretic spatial model. Our results demonstrate that combining adaptive therapy with migrastatics effectively delays the onset of metastasis and reduces both the number and size of metastases across most cancer scenarios analyzed. This approach not only extends the time to the first metastasis but also enhances the overall efficacy of adaptive therapies. Our findings suggest a promising new direction for cancer treatment, where adaptive therapy, in combination with migrastatic agents, can target both the evolution of resistance and the metastatic spread of cancer cells.

## Introduction & Motivation

Cancer represents the second leading cause of death worldwide. Moreover, recent trends show that cancer may overtake cardiovascular disease by approximately 2030, becoming the first leading cause of death [15, 17]. For patients with cancer, treatment is commonly aimed at complete eradication of the tumor, i.e. targeting the uncontrolled proliferation of the cancer cells [46, 49, 16].

In standard of care, patients are typically given the maximum tolerated dose (MTD), which is the highest dose that a mean patient can handle without experiencing intolerable toxicity [52, 49, 2]. While MTD-based therapy offers survival benefits, it often comes with severe side effects. Moreover, recurrence is almost inevitable due to the emergence of therapeutic resistance [38, 51, 23, 32, 20]. To address these challenges, adaptive therapy (AT), also known as evolutionary therapy, has been proposed [50, 27, 3, 10, 25]. AT involves administering therapy doses based on the current state of tumor growth and anticipated evolutionary changes and hence, mathematical models have been crucial in its development [1, 21, 12, 10, 16]. The goal of adaptive therapy is often to maintain a sufficient population of drug-sensitive cells, enabling them to outcompete drug-resistant cells, and hereby control the tumor the tumor burden longer. A pilot example of such a therapy was the one proposed by Zhang et al. in metastatic castrate-resistant prostate cancer (mCRPC) [43, 13]. In this clinical trial, mCRPC has been treated with MTD until the total tumor burden is halved compared to its initial size. Upon reaching this threshold, the treatment is stopped until the tumor burden reaches its original size, allowing for competition between drug-sensitive and resistant cells. When the total tumor burden reaches its initial size, treatment with MTD is reinstated and a new treatment cycle begins. Thus, while complete tumor eradication may not be possible, Zhang et al’s protocol aims to control the tumor burden through limiting the development of uncontrollable drug resistance.

While paradigm shift towards adaptive therapy is already happening, most existing adaptive therapy protocols aim at tumor containment, control or eradication [43, 24, 21, 10, 1], and they do not target spread of the disease. However, up to 90% of mortality in solid tumors is due to metastasizing rather than cancer growth alone [48, 28]. In order for the primary tumor to successfully metastasize, cancer cells need to complete a number of sequential events, the so-called invasion-metastatic cascade [45]. Firstly, some cancer cells acquire an invasive phenotype that allows them to dissociate from the original tumor mass and invade the surrounding stroma. Cancer cells then intravasate into the bloodstream, where they have to withstand strong selective pressure and unfavorable conditions, extravasate into the premetastatic niche and finally form a secondary lesion [41].

Current systemic approaches, including screening, chemotherapy, targeted therapy and immunotherapy, aimed at inhibiting tumor growth, ignore the invasive potential of cancer cells, and some may even support metastasis forming [30]. Therefore, metastasis may be responsible for the majority of therapy failures since no efficient therapeutic strategies currently target the metastatic cascade, making metastatic cancer highly incurable and fatal.

Recently, migrastatics, i.e. drugs targeting cancer cell motility and invasion, have been proposed as a potentially more rational therapeutic approach [39, 8, 5]. Migrastatics refer to a specific class of drugs that interfere with various mechanisms involved in cancer cell invasion and, therefore, with its ability to metastasize. Unlike conventional cytostatic drugs that primarily target cell proliferation, migrastatics focus on inhibiting local invasion and metastasis. Hereby they address the motility and invasive aspect of cancer and thereby prevent the progression of primary tumor into metastatic cancer.

Moreover, migrastatic strategies possess some important advantages compared to commonly used cytotoxic therapies. Even though the cytotoxic therapy may result in the shrinkage of the tumor, migration capacity and the metastatic potential of the resistant cells are not affected. Therefore, these cells retain the capability of forming secondary loci in distant organs. Contrarily, implementing the migrastatic approach does not lead to the shrinkage of the tumor, because these drugs are neither cytotoxic nor antiproliferative. However, the motility of cancer cells within the tumor microenvironment is significantly reduced. One of the biggest advantages of a successful migrastatic therapy would lie within the possible decrease of currently used high-dose cytotoxic treatment.

Conventional cytotoxic and/or cytostatic therapeutics target uncontrollable proliferation, which ultimately results in Darwinian selection of resistant clones within the tumor mass [31]. The indisputable advantage of migrastatic drugs would lie in their ability to inhibit cancer cell motility and invasion without disrupting the proliferative signaling. Therefore, in case that some cancer cells would become resistant to migrastatic therapy, they would not gain a proliferative advantage and would not get enriched within the cancer cell population, making metastasizing much less effective.

Migrastatics are novel drugs that do not yet have a defined administration regime, and since their effect is distinct from that of cytostatics, it is crucial to adopt a new approach for their administration. Three specific regimens of therapeutic use of migrastatic drugs have been proposed to combat the formation and progression of metastases [5]. Firstly, the neoadjuvant/adjuvant therapy suggests the administration of migrastatics before and after surgical procedures. The inclusion of migrastatics seeks to counteract and/or to minimize the risk of tumor cells initiating a metastatic program as a result of pro-invasive changes in their environment caused by wound healing processes (pro-migratory effect of cytokines) and post-operative treatments (anticoagulants) [33]. Secondly, the combination of cytostatics and migrastatics drugs is proposed as a means to effectively reduce the development of metastases. Combining these two types of treatments aims to target both the primary tumor and the migratory potential of cancer cells, thereby reducing the likelihood of metastatic spread. Lastly, the migrastatic therapy aimed at minimizing the long-term risk of metastasis. It can be used alone to slow down or even prevent metastasis, thereby reducing the tumor burden. Alternatively, in combination with either non-systemic treatment targeting only the primary tumor, such as surgery or radiotherapy, or in combination with immunotherapy.

We believe that including migrastatics into both classical and adaptive treatment protocols has a high potential for both controlling cancer growth and avoiding tumor metastases at the same time. To illustrate the potential of including migrastatics into the existing treatment protocols, here we present a spatial game-theoretic model similar to that of You et al. (2017) [42] that allows us to examine a potential impact of applying both cytostatic and migrastatic treatments and that of applying migrastatic treatment only. As cytostatic treatment, we investigate both the standard of care and adaptive treatment approach to evaluate whether combining current adaptive protocols with migrastatics would be beneficial.

## Methods

To examine the potential of migrastatics in the standard of care and metastatic treatments, we have developed a continuous-space evolutionary game-theoretic model of metastatic cancer growth and its spread to potential metastatic sites. This model allows us to explore the impact of cytostatic and migrastatic treatments on the tumor growth, formation of metastases, and composition of these metastases. The model includes the primary tumor and eight possible migration sites. The cancer cells are either sensitive or resistant to the cytostatic treatment and may migrate and potentially form metastases. Our model includes both frequency-dependent and density-dependent selection, as cancer cells’ probability of producing daughter cells is given by their pairwise interaction with other cells within their neighborhood and is captured in the fitness matrix. Different fitness matrices correspond to different scenarios regarding the ability of sensitive cancer cells to outcompete the resistant ones with and without treatment (see [**Rapoport1966Taxonomy**, **Hofbauer1998Evolutionary**, **Skyrms2004StagHunt**] for classificaiton of the possible matrices, including coordination, anti-coordination, harmony, prisoner’s dilemma). We select specific examples of such matrices for our cases studies to demonstrate the qualitative results corresponding to specific class of fitness matrices under different forms of treatment.

### Model dynamics

The model contains one primary tumor site with 8 possible migration sites and has infinite continuous space. The field is defined as a square [−*L, L*] × [−*L, L*] where *L* is large. The primary site is placed in the centre of the field and is defined as a disc of a prespecified radius with a centre at (0, 0). We assume that metastases can potentially form at 8 metastasis sites, which are located at discs with a predefined radius equal to the interaction radius and placed in the field with their center the locations (−60, 60), (0, 60), (60, 60), (−60, 0), (60, 0), (−60, 60), (0, −60), (60, −60).

At the start of the simulation, a fixed number of cells with a predefined fraction of cells sensitive (type *S*) and resistant (type *R*) to cytostatic treatment are randomly placed within the primary site. Of these cells a pre-specified fraction is selected as invasive and have the potential to migrate to one of the eight migration sites, or from one of the migration sites to the primary tumor at later generations. The same starting configuration, i.e. the placement of the cells of primary tumor, is used for all case studies.

Interactions happen in generations, similarly to [42]. Each simulation is simulated for 100 generations. Within each generation, each cell in the field is selected as a focal cell at random order and undergoes the following steps:

1. *Death*: The focal cell may die according to a pre-specified death probability, equal for all cells in the simulation.
2. *Migration*: If the focal cell survives and has the property to be invasive, with a pre-specified probability the cell migrates to a migration site that is randomly selected from the eight possible migration sites. If the cell migrates, the cell then survives at the new site with a pre-specified survival probability.
3. *Proliferation*: In the last part of one generation the cells can proliferate, hereto an interaction partner is randomly selected from the focal cell’s interaction neighbourhood, defined as a disc with center at the focal cell and radius given by the interaction radius. The proliferation happens only if the number of cells within the focal cell’s neighborghood is below the predefined carrying capacity. The interaction determines the probability of the focal cell to create an offspring of its own type, accumulated in the fitness matrix *A* = (*a_ij_*)_2_*_×_*_2_. For a focal cell of type *i* and an interaction partner of type *j*, where *i, j* ∈ {*S, R*}, element *a_ij_* of the fitness matrix *A* defines the probability that the focal cell produces a daughter cell of its own type *i*, which is then placed randomly in the interaction neighborhood. Daughter cells in the current generation cannot be chosen as interaction partners until the next generation, but are taken into account when evaluating whether the carrying capacity has not been reached.

### Case studies

We investigate impact of different treatment scheduels for our model with 5 different fitness matrices, each corresponding to different interactions between the sensitive and resistant cancer cells within the model.

Three of these matrices represent competition between sensitive and resistant cell populations are evaluated. In these three matrices interactions of a cell with its own type result in a probability of proliferation of 0.2 for both sensitive and resistant cell types. Where proliferation probabilities when interacting with a partner of a different type are varied between matrices.

The first matrix is the anti-coordination sensitive *(AC-S)* matrix [7]:

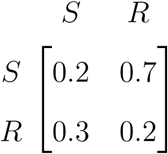

Here the sensitive cells have a higher probability to proliferate when they interact with resistant cells than when they interact with cells of its own type. Resistant cells have a higher chance to proliferate when interacting with sensitive cells than with their own type. In the mostly resistant environment, sensitive cells will grow more than sensitive cells, while in the mostly sensitive environment, resistant cells will grow more than in the sensitive cells.

The second matrix, anti-coordination resistant *(AC-R)*, switches the roles of the sensitive and resistant cells [7]:

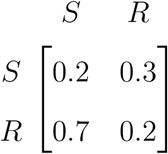

The third matrix, anti-coordination equal *(AC-E)*, is a symmetric anti-coordination game [7], defined by the following fitness matrix:

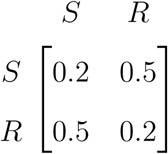

We will define to this game as *Competition equal* in our case studies.

Cancer cells of the same type could also cooperate with each other during tumor growth. Here we evaluate two cases, one where sensitive cells benefit from interactions with their own type through the highest proliferation chance (indeed “S” is an evolutionarily stable strategy here), while all other interactions bring a lower proliferation chance, and a matrix where the same holds for the resistant cells (with “R” being the evolutonarily stable strategy). These games, defined via fitness matrices

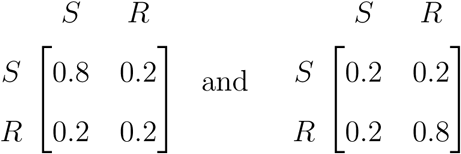

will be referred to as *Cooperation sensitive* and *Cooperation resistant*, respectively.

Note that these two games are at the transition between harmony and prisoner’s dilemma games [7].

For each of the five fitness matrices four treatment strategies will compared to each other and to no treatment, in terms of how long the tumor burden can be contained. The four treatment strategies combine two treatments:

1. *Cytostatic treatment* : When cystostatic treatment is applied a predefined percentage of sensitive cells are killed and removed immediately. Resistant cells are not affected.
2. *Migrastatic treatment* : When migrastatic treatment is applied the probability of migration is lowered ten times.

We compare (i) applying migrastatics only, (ii) no treatment, (iii) cytostatic MTD treatment without migrastatics, (iv) cytostatic MTD treatment with migrastatics, (v) adaptive cytostatic treatment (AT) without migrastatics, and (vi) adaptive cytostatic treatment with migrastatics. Here AT refers to Zhang et al.’s protocol [43, 14].

All parameter values are shown in Appendix A.

## Results

### Migrastatics prevent metastases and tumor growth

Firstly, we evaluate the effect of migrastatics alone on the case studies with the 5 fitness matrices. Figure 1 summarizes the result of the case study with fitness matrix *AC-S*, while Annex B summarizes case studies with the other matrices. In all case studies, adding migrastatics slows down the increase of total tumor burden when compared to no treatment (Figure 1A). This can potentially be due to the limitation of space when the probability of migration is lowered. As expected, with migrastatics, the number of metastases is lower, while time to the first metastasis is longer (Figure 1B). In addition to lower total tumor burden and later metastasizing, the formed metastases contain less cells (Figure 1C).

**Figure 1:**
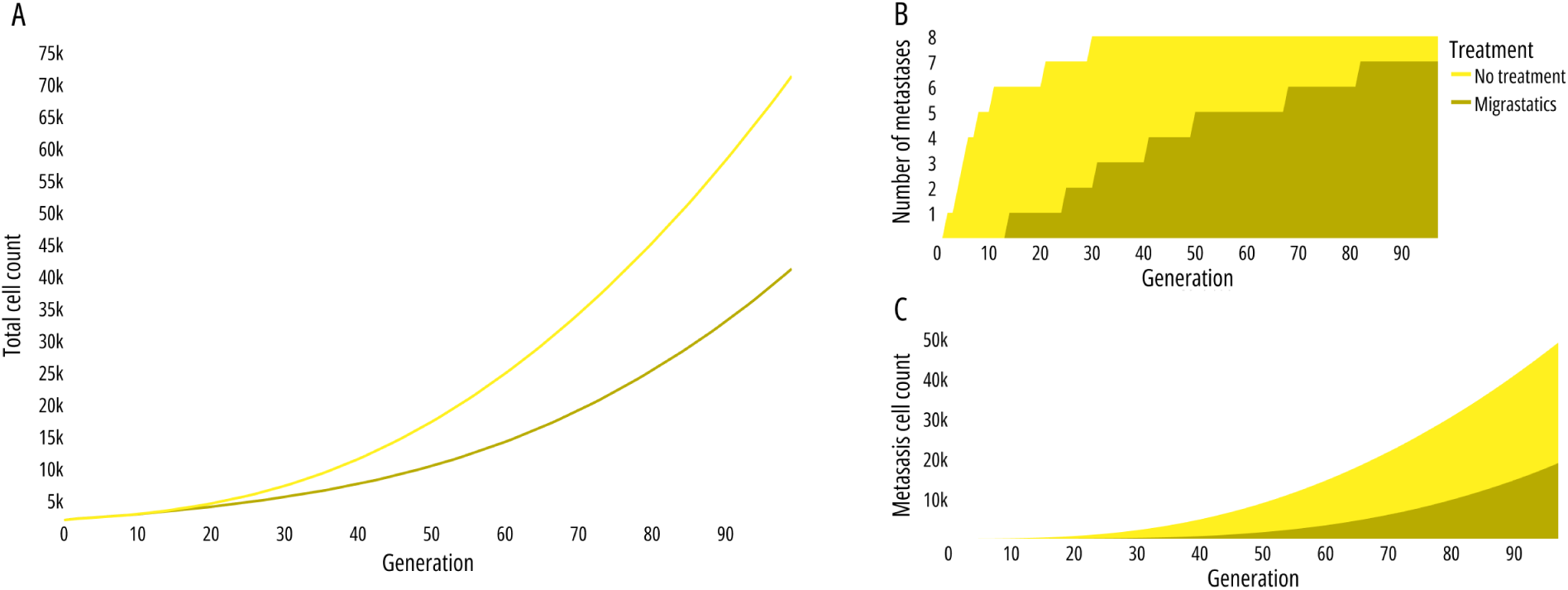
Dynamics of cancer cells under migrastatic treatment for the case study with the *AC-S* fitness matrix: Migrastatic treatment reduces total tumor burden (A) by decreasing metastasis formation (B) and size of metastases (C). Time to first metastasis increases, and average metastasis size decreases. Results are averaged over 50 simulations.

**Figure 2:**
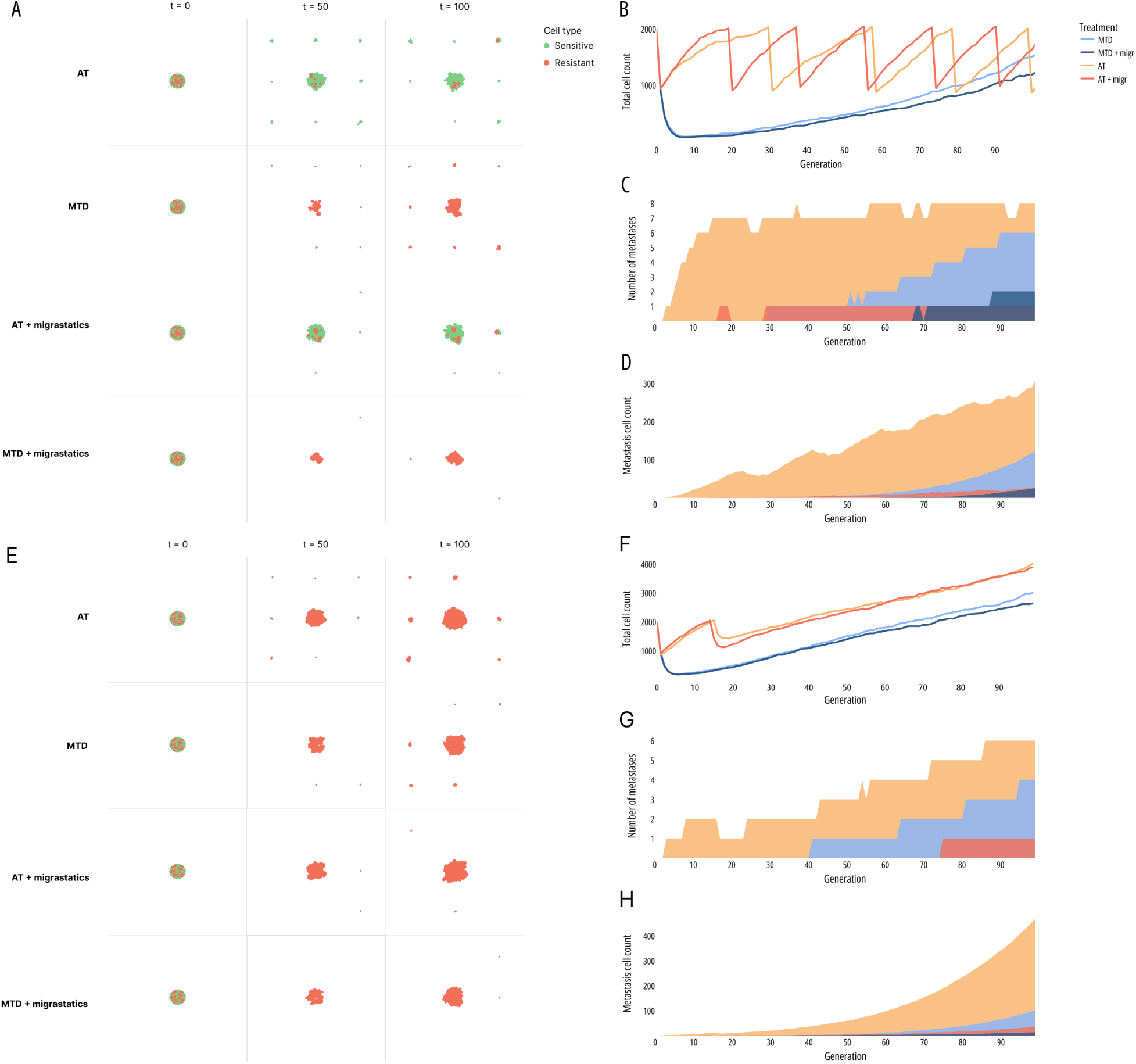
Cancer cell growth under the *cooperation sensitive* (A-D) and *resistant* (E-G) cases with four treatments combining Adaptive Therapy (AT) or Maximum Tolerable Dose (MTD) with and without migrastatics. **(A)** & **(E)** Reduced metastases and persistence of sensitive cells with AT **(A)**. **(B)** & **(F)** Migrastatics do not inhibit overall growth. **(F)** AT fails with tumors becoming fully resistant. **(C)** & **(G)** Delayed and reduced metastases, especially with AT. **(D)** & **(H)** Smaller, delayed metastases with migrastatics. **(C)**, **(D)**, **(G)** and **(H)** averaged over 50 runs.

### Combining adaptive therapy and migrastatics hinders metastasizing

Current cancer treatment is not geared towards preventing metastases, but towards killing cancer cells. Current standard of care is treatment with MTD, however AT has shown great potential in increasing time to progression [43, 13]. Thereto we analyze the effect of migrastatics combined with cystostatic treatments.

Firstly, we note that the addition of migrastatics to Zhang et al’s adaptive therapy protocoldoes not alter its oscillatory behavior in all evaluated cases (Figure 3 - 5B & F).

**Figure 3:**
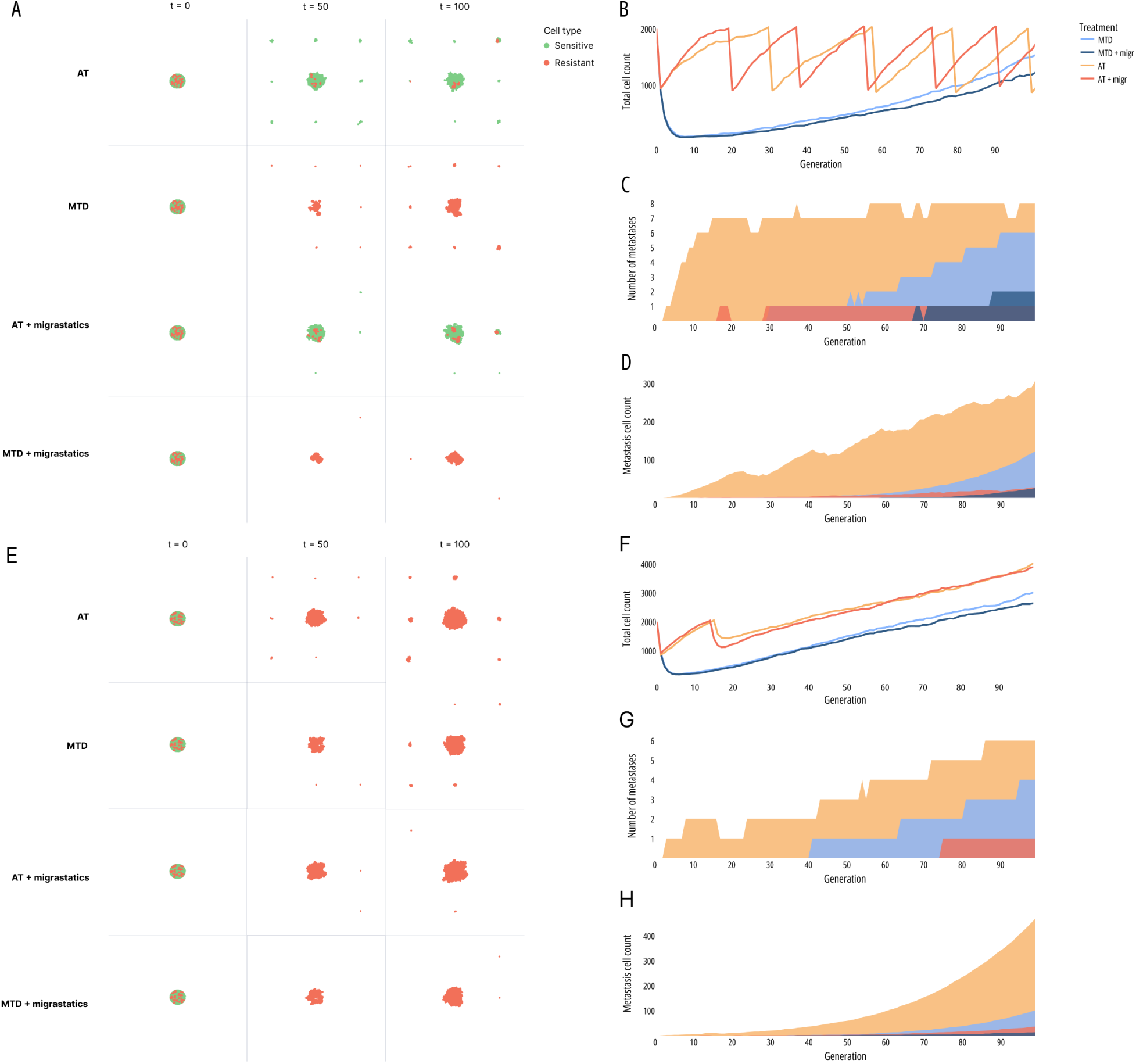
Cancer cell growth under the *cooperation sensitive* (A-D) and *resistant* (E-G) cases with four treatments combining Adaptive Therapy (AT) or Maximum Tolerable Dose (MTD) with and without migrastatics. **(A)** & **(E)** Reduced metastases and persistence of sensitive cells with AT **(A)**. **(B)** & **(F)** Migrastatics do not inhibit overall growth. **(F)** AT fails with fully resistant tumors. **(C)** & **(G)** Delayed and reduced metastases, especially with AT. **(D)** & **(H)** Smaller, delayed metastases with migrastatics. **(C)**, **(D)**, **(G)** and **(H)** averaged over 50 runs.

In all but one of the cases AT prolongs time to progression, as expected. In the case of *Cooperation resistant* case, AT becomes ineffective around the 20th generation, due to all sensitive cells being killed by the cytotoxic drugs (Figure 3F). After this generation, the increase of the total tumor burden per generation then becomes similar for MTD and AT and because the cell population at that generation is larger at this generation in the AT conditions the outcome of the total tumor burden is higher is these conditions.

In the *Cooperation sensitive* case, AT still facilitates oscillations at the 100th generation (Figure 3B), and cells sensitive to the cytotoxic treatment are more frequent than resistant cells at this generation (Figure 3A). Similarly to the *Cooperation resistant* case, the addition of migrastatics to the cytotoxic treatment does prolong time to the first metastases. Metastases occur first in conditions treated with AT, followed by MTD and last in MTD combined with migrastatics (if at all). With both cooperative fitness matrices, the average number of cells per metastasis when migrastatics are applied is low. In the *Cooperation sensitive* case, metastases occur earlier when migrastatics are combined with AT than when they are combined with MTD. However, at the 100th generation, there are no sensitive cells present for MTD with migrastatics, while they are still there when AT and migrastatics are combined.

In the *AC-E* case, only resistant cells remain at generation 50 (Figure 4A) and cycling of AT has stopped (Figure 4B). In the other two competition cases, *AC-S* and *AC-R*, the application of AT does prolong time to progression, as sensitive cells are still present at generation 50, with total tumor burden still oscillating at generation 100 for the *AC-S* case (Figure 5A). In all competition cases, migrastatics prolongs time to metastasizing, and average metastasis size is smaller at the 100th generation for all cases, with best results for AT combined with migrastatics.

**Figure 4:**
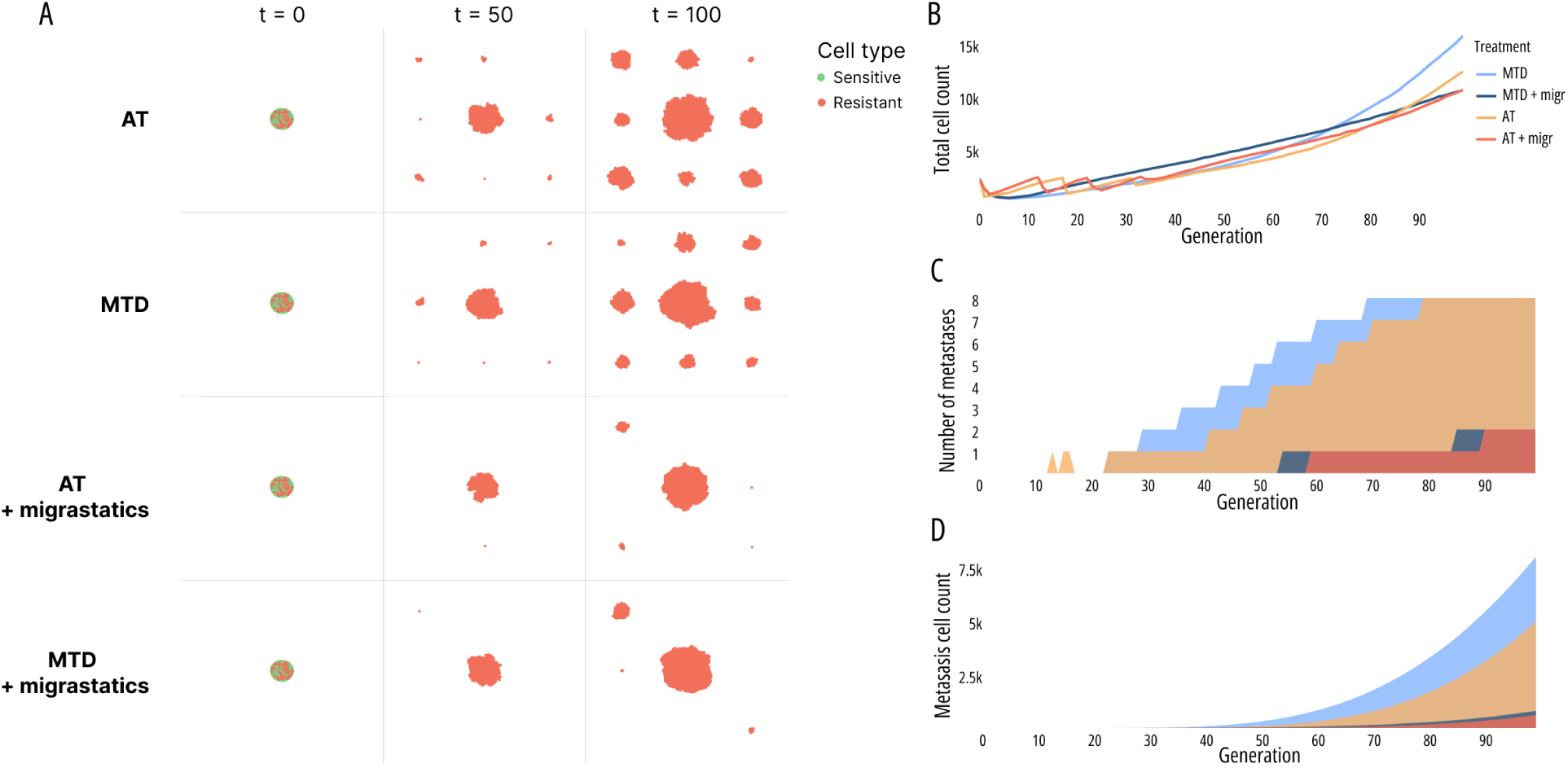
Cancer cell growth under the *AC-E* fitness matrix with treatments combining Adaptive Therapy (AT) or Maximum Tolerable Dose (MTD) with and without migrastatics. **(A)** Reduced metastases with migrastatics. **(B)** Migrastatics do not inhibit overall cancer growth; AT fails with completely resistant tumors. **(C)** Delayed and reduced metastases with migrastatics. **(D)** Smaller, delayed metastases with migrastatics **(C)** and **(D)** averaged over 50 runs.

**Figure 5:**
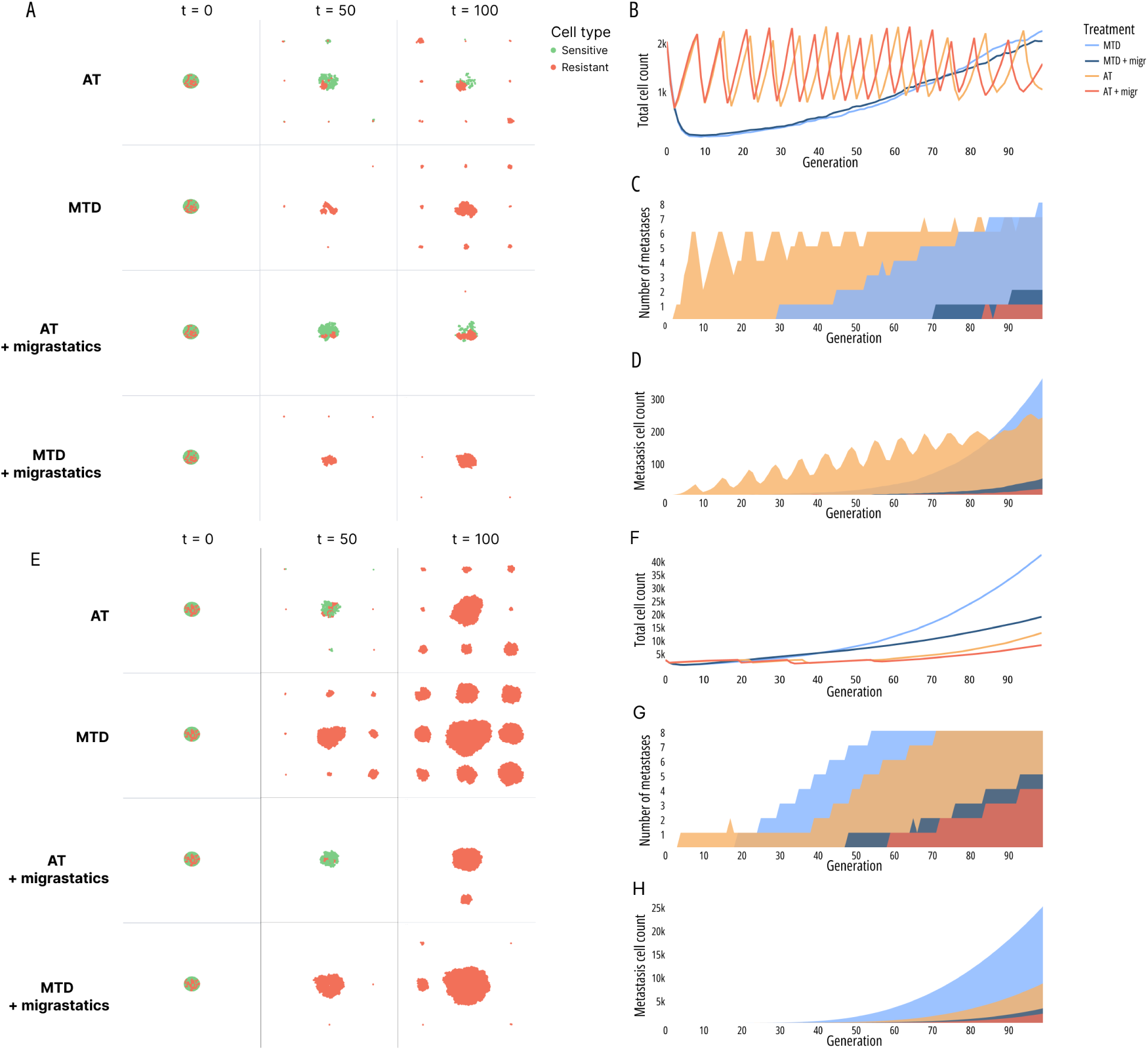
Cancer cell growth under the *AC-S* (A-D) and *AC-R* (E-H) fitness matrices with treatments combining Adaptive Therapy (AT) or Maximum Tolerable Dose (MTD) with and without migrastatics: **(A)** AT preserves sensitive cells; MTD leads to resistance. With both treatment strategies migrastatic treatment reduces metastases. **(E)** Reduced metastases with migrastatics. **(B)** & **(F)** Migrastatics do not affect overall cancer growth. **(F)** AT fails after its three cycles, due to resistance. **(C)** & **(G)** Migrastatics delay and reduce metastasis; metastases occur earlier with AT due to higher cell numbers. **(D)** & **(H)** Smaller, delayed metastases with migrastatics. **(C)**, **(D)**, **(G)** and **(H)** averaged over 50 runs.

**Figure 6:**
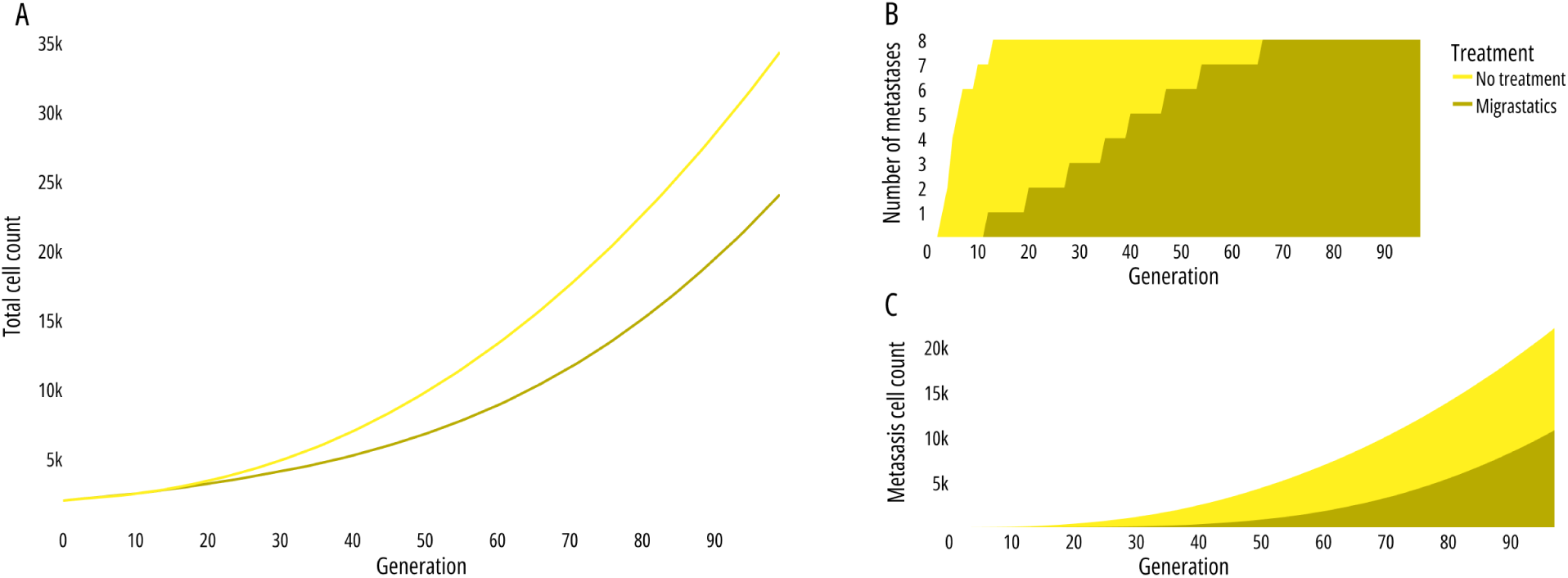
Growth of cancer cells for the pay-off matrix for *Anti-coordination equal* under no treatment versus migrastatic treatment. **(A)** Migrastatic treatment significantly reduces total tumor burden over 100 generations. **(B)** This is attributed to delayed and diminished metastasis formation, with metastases arising later. **(C)** Moreover, the average metastasis size remains consistently lower with migrastatic treatment. All results averaged over 50 simulations.

## Discussion

The aim of this study was to identify a treatment strategy that does not only focus on the inhibition of tumor growth while limiting the number of resistant cancer cells, but additionally reduces the formation of metastases. This was to demonstrate a high potential that migrastatic treatment may have, especially in combination with adaptive therapy treatments.

We have demonstrated that Zhang et al’s adaptive therapy protocol [43, 13] is successful in postponing time to progression when compared to the standard of care for most scenarios studied. This is because this protocol prevents outgrow of resistant cell types of the tumor early after treatment has started, successfully facilitating the survival of drug-sensitive cancer cell populations. However, due to the high total tumor burden the Zhang et al’s maintains, it promotes cancer cell migration and forming metastases. This often results in earlier metastasizing of the tumor and larger metastases. These results point to the need to expand the scope of adaptive therapies beyond drug resistance alone and include targetting cancer’s invasiveness.

When migrastatics are added to both the MTD and AT treatment protocols, we observe both a decrease in the number and size of metastases and an increase in the time to metastasis for all cases evaluated here. For fitness matrices where AT prolonged time to treatment failure, addition of migrastatics to the AT improved the time to progression, number of metastases, and time to the first metastasis. Together, our modelling results demonstrate that the combination of migrastatics with AT can help target both tumor resistance and metastasizing. Future work will be needed to validate these findings in *in vitro* and *in vivo* studies.

While there is implicitly no reason for prevailing of cells which would eventually get resistant to migrastatics, because such resistance would not give them a proliferative advantage [5, 40, 31], the synergistic effect with AT will have yet to be validated in preclinical and clinical studies. Initially, the combination of AT with migrastatic therapy will be tested in 3D spheroid model [44] and subsequently in a suitable mouse model of metastasis, tailored for testing of migrastatic drug efficacy, e.g. [18]. Data collected during these experimental studies can be used to validate our models and optimize adaptive therapy enhanced by migrastatics.

Our model assumes a fixed location of the primary site and potential metastatic sites. Future studies should validate our results when assuming varying potential number of metastases as well as varying locations of both the primary site and potential metastatic sites. This is because the location of the primary tumor is a determining factor to where metastases will form and the metastasis location in itself are correlated with patient survival [37, 26]. An assumption presented here is that the cells in the cancer model are fully sensitive or resistant [35], while the resistance induced by treatment of cancer cells can also evolve in the direction of the fitness gradient [36, 47, 19, 12]. Future work could include quantitative resistance or a combination of qualitative and quantitative resistance, and study effect of such assumptions on the success of treatment protocols we have described here.

The interactions values of the fitness matrices used here are chosen to describe different scenarios with different possible conditions for sensitive and resistant cells and demonstrate the effect different treatments in such scenarios. These results show that AT can be successfully applied in cases of an anti-coordination game. However, with the *Cooperation resistant* matrix, where resistant cells have a high benefit from interacting with their own type, AT does not improve time to progression when compared to MTD. Here the resistant cells outcompete the sensitive cells and the tumors become fully resistant, causing earlier treatment failure due to the higher total tumor burden. However, even when AT is ineffective, the results of the *Cooperation resistant* case demonstrate that adding migrastatics to the administered treatment is still effective in reducing metastases. Moreover, one can consider other forms of adaptive therapy than Zhang et al’s protocol, such as double-bind or extinction therapies [29, 3], which will likely be more effective and can still be combined with migrastatics. Treatment optimization to choose the best treatment combination and sequence using optimal control theory is also a potential next treatment step [34, 22, 4].

We have demonstrated that by combining Zhang et al’s AT protocol [43] with migrastatics both resistance and metastasis can be controlled longer and treatment failure is hereby delayed. This points to a promising new treatment strategy combating both the proliferative as the invasive aspects of cancer. Future work will need to focus on extending the current model and validating these findings with *in vivo* and *in vitro* data, similarly as it was done for other mathematical models [**kazna**, 6, 11, 9]

## Funding

This research was supported by European Union’s Horizon 2020 research and innovation program under the Marie Sk-lodowska-Curie grant agreement No 955708 and the Dutch Research Council projects OCENW.KLEIN.277 and VI.Vidi.213.139.

## Annex A: Parameter values for our model

Parameter values from Table 1 are common for all scenarios when no treatment is applied.

**Table 1:** Parameters common for all scenarios when neither cytostatic nor migrastatic treatment is applied.

| Description | Value |
| --- | --- |
| Number simulation runs | 50 |
| Initial type distribution | 97.5% sensitive cells, 2.5% resistant cells |
| Carrying capacity (cells per unit area) | 6 |
| Natural death probability | 0.15 (Anti-Coordination) 0.10 (Cooperation) |
| Density radius | 1 |
| Reproduction radius | 1 |
| Interaction radius | 1 |
| Number migration sites | 8 |
| Survival probability at new site | 0.1 |
| Fraction of invasive cells | 0.1 |
| Migration probability | 0.1 |
| Migration probability with migrastatics | 0.01 |

Furthermore under treatment, the parameters from Table 2 are applied.

**Table 2:** Parameters applied under treatment.

|  |  |
| --- | --- |
| Fraction of sensitive cells killed under cytostatic treatment | 0.6 |
| Probability to migrate under migrastatic treatment | 0.01 |

## Annex B: Supplementary results

**Figure 7:**
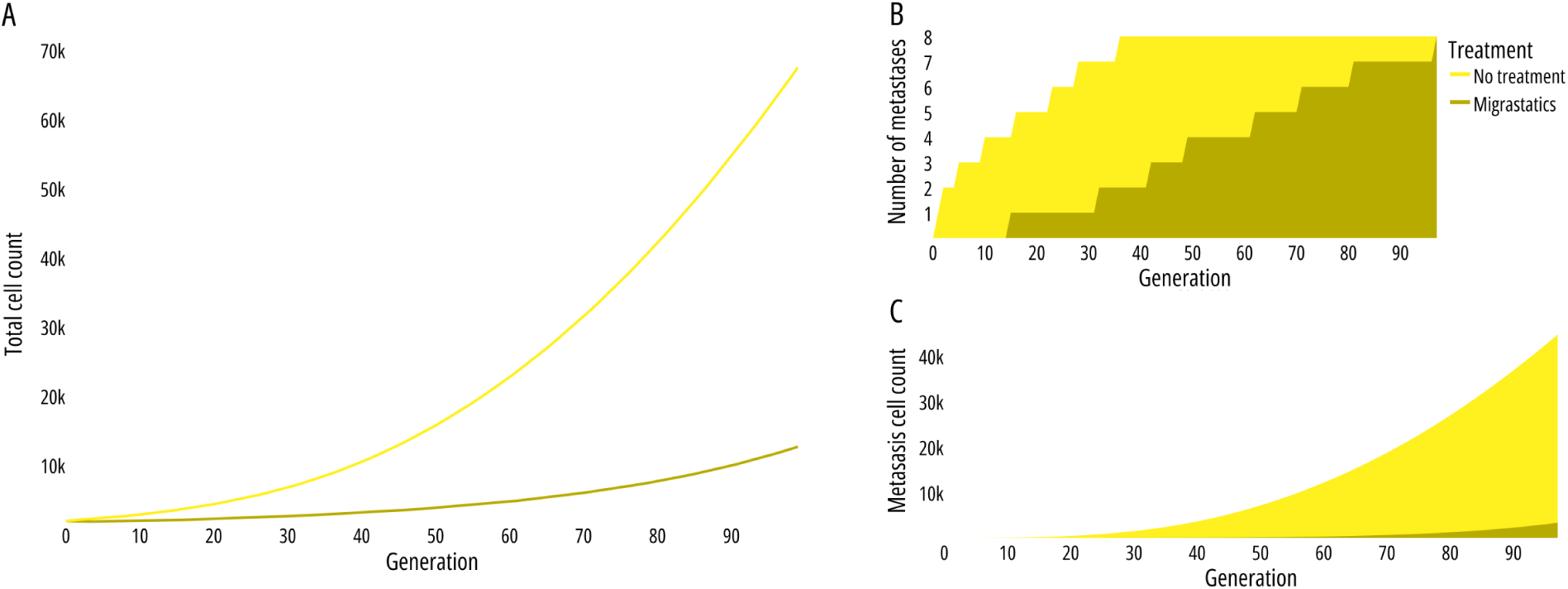
Growth of cancer cells for the *Anti-coordination* fitness matrix under no treatment versus migrastatic treatment: **(A)** Over 100 generations, migrastatics consistently reduce overall tumor burden. **(B)** This effect stems from delayed and reduced metastasis formation, with metastases emerging later. **(C)** Migrastatic treatment consistently maintains smaller metastasis size. All results are averaged over 50 simulations.

**Figure 8:**
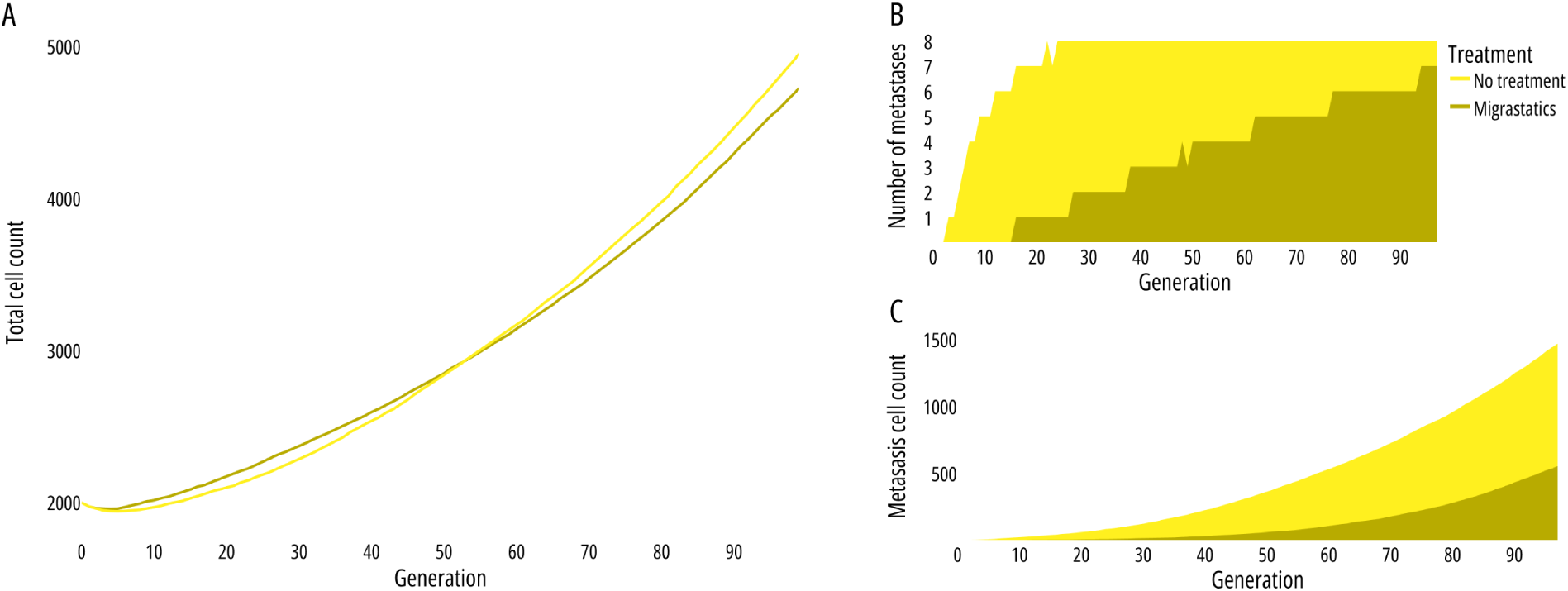
Growth of cancer cells for the *Cooperation sensitive* fitness matrix under no treatment versus migrastatic treatment: **(A)** Migrastatics effectively reduce overall tumor burden over 100 generations. **(B)** This is primarily due to a lower metastasis incidence, with metastases forming later. **(C)** Migrastatic treatment results in smaller metastasis sizes. All results are averaged over 50 simulations.

**Figure 9:**
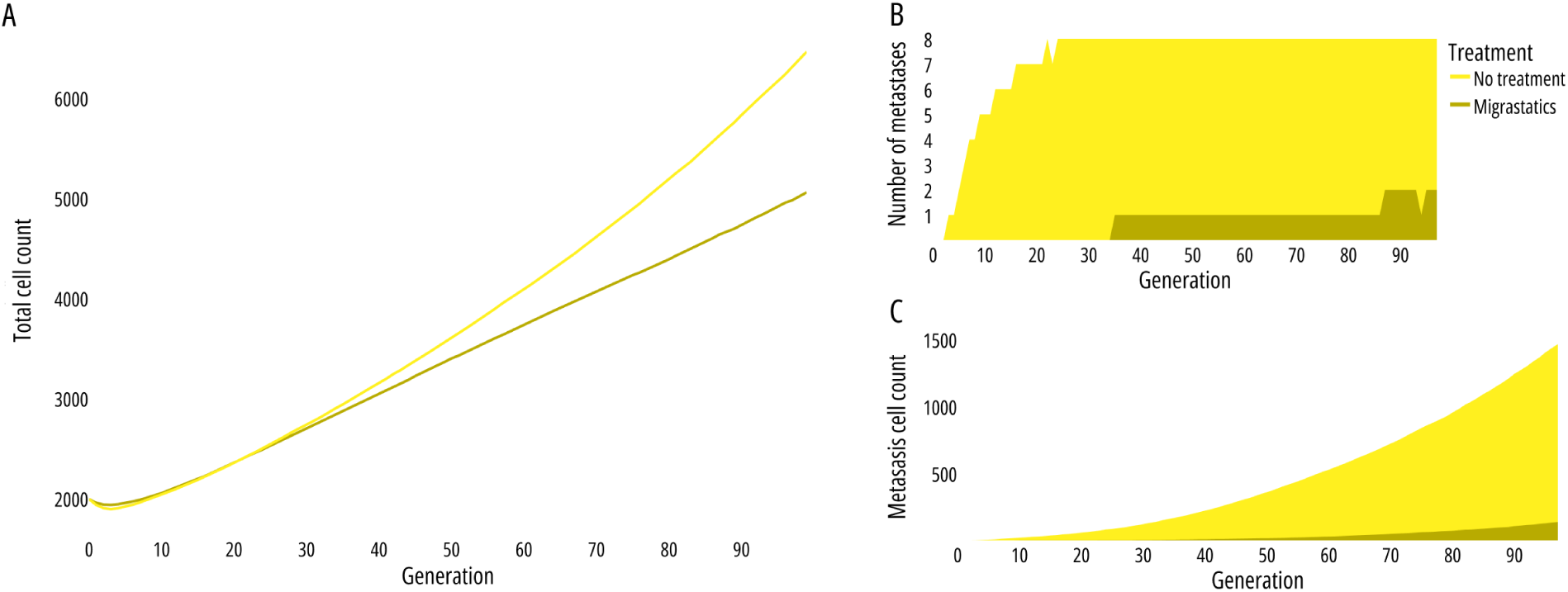
Growth of cancer cells for the *Cooperation resistant* fitness matrix under no treatment versus migrastatic treatment. **(A)** Migrastatic treatment effectively reduces overall tumor burden over 100 generations. **(B)** This effect stems from delayed and reduced metastasis formation with metastases emerging later. **(C)** Migrastatic treatment consistently maintains smaller metastasis size. All results are averaged over 50 simulations.

## Notes

### Competing Interest Statement

The authors have declared no competing interest.

## References

[1] Alvarez, Frank Ernesto and Viossat, Yannick. “Tumor containment: a more general mathematical analysis”. In: Journal of Mathematical Biology 88.4 (2024), p. 41.

[2] Gad, Shayne C. “Maximum tolerated dose”. In: Encyclopedia of Toxicology. Vol. 6. Elsevier, Jan. 2024, pp. 43–44. doi: 10.1016/B978-0-12-824315-2.00532-7.

[3] Luddy, Kimberly A, et al. “Evolutionary double-bind treatment using radiotherapy and NK cell-based immunotherapy in prostate cancer”. In: bioRxiv (2024), pp. 2024–03.

[4] Salvioli, Monica et al. “Stackelberg evolutionary games of cancer treatment: What treatment strategy to choose if cancer can be stabilized?” In: Dynamic Games and Applications (2024). doi: 10.1007/s13235-024-00609-z.

[5] Škarková, Aneta, et al. “Educate, not kill: treating cancer without triggering its defenses”. In: Trends in molecular medicine 30.7 (July 2024), pp. 673–685. issn: 1471-499X. doi: 10.1016/J.MOLMED.2024.04.003. url: https://pubmed.ncbi.nlm.nih.gov/38658206/.

[6] Soboleva, Arina, et al. “Validation of polymorphic Gompertzian model of cancer through in vitro and in vivo data”. In: PLOS ONE, in print (2024).

[7] Bayer, Péter and West, Jeffrey. “Games and the Treatment Convexity of Cancer”. In: Dynamic Games and Applications 13.4 (Dec. 2023), pp. 1088–1105. issn: 21530793. doi: 10.1007/S13235-023-00520-Z/FIGURES/7. url: https://link.springer.com/article/10.1007/s13235-023-00520-z.

[8] Raudenská, Martina, et al. “Engine shutdown: migrastatic strategies and prevention of metastases”. In: Trends in Cancer 9.4 (Apr. 2023), pp. 293–308. issn: 2405-8033. doi: 10.1016/J.TRECAN.2023.01.001.

[9] Strobl, Maximilian et al. “Adaptive therapy for ovarian cancer: An integrated approach to PARP inhibitor scheduling”. In: bioRxiv (Mar. 2023). Preprint. doi: 10.1101/2023.03.22.533721. url: https://www.biorxiv.org/content/10.1101/2023.03.22.533721v1.

[10] West, Jeffrey et al. “A survey of open questions in adaptive therapy: Bridging mathematics and clinical translation”. In: eLife 12 (2023), e84263. doi: 10.7554/eLife.84263.

[11] Ghaffari Laleh, N. et al. “Classical mathematical models for prediction of response to chemotherapy and immunotherapy”. In: PLOS Computational Biology 18.2 (Feb. 2022), pp. 1–18.

[12] Wölfl, Benjamin, et al. “The contribution of evolutionary game theory to understanding and treating cancer”. In: Dynamic Games and Applications 12.2 (2022), pp. 313–342.

[13] Zhang, Jingsong et al. “Evolution-based mathematical models significantly prolong response to abiraterone in metastatic castrate-resistant prostate cancer and identify strategies to further improve outcomes”. In: eLife 11 (June 2022). issn: 2050-084X. doi: 10.7554/eLife.76284.

[14] Zhang, Jingsong et al. “Evolution-based mathematical models significantly prolong response to abiraterone in metastatic castrate-resistant prostate cancer and identify strategies to further improve outcomes”. In: eLife 11 (June 2022). issn: 2050084X. doi: 10.7554/ELIFE.76284.

[15] Bray, Freddie et al. “The ever-increasing importance of cancer as a leading cause of premature death worldwide”. In: Cancer 127.16 (Aug. 2021), pp. 3029–3030. issn: 1097-0142. doi: 10.1002/CNCR.33587.

[16] Dujon, Antoine M., et al. “Identifying key questions in the ecology and evolution of cancer”. In: Evolutionary applications 14.4 (2021), pp. 877–892.

[17] Ganesh, Karuna and Massagué, Joan. “Targeting metastatic cancer”. In: Nature Medicine 2021 27:1 27.1 (Jan. 2021), pp. 34–44. issn: 1546-170X. doi: 10.1038/S41591-020-01195-4. url: https://www-nature-com.utrechtuniversity.idm.oclc.org/articles/s41591-020-01195-4.

[18] Maiques, Oscar et al. “A preclinical pipeline to evaluate migrastatics as therapeutic agents in metastatic melanoma”. In: British journal of cancer 125.5 (Aug. 2021), pp. 699–713. issn: 1532-1827. doi: 10.1038/S41416-021-01442-6. url: https://pubmed.ncbi.nlm.nih.gov/34172930/.

[19] Pressley, M., et al. “Evolutionary dynamics of treatment-induced resistance in cancer informs understanding of rapid evolution in natural systems”. In: Frontiers in Ecology and Evolution 9 (2021), p. 460.

[20] Strobl, M., et al. “Turnover modulates the need for a cost of resistance in adaptive therapy”. In: Cancer Research 81.4 (2021), pp. 1135–1147.

[21] Viossat, Y. and Noble, R. “A theoretical analysis of tumour containment”. In: Nature Ecology and Evolution (2021).

[22] Cunningham, Jessica J, et al. “Optimal Control to Reach Eco-Evolutionary Stability in Metastatic Castrate Resistant Prostate Cancer”. In: PLOS ONE 15.12 (2020), pp. 1–24.

[23] Dinić, Jelena, et al. “Repurposing old drugs to fight multidrug resistant cancers”. In: Drug Resistance Updates (2020), p. 100713.

[24] Gatenby, R. A., et al. “Eradicating metastatic cancer and the eco-evolutionary dynamics of Anthropocene extinctions”. In: Cancer research 80.3 (2020), pp. 613–623.

[25] Gluzman, Mark, Scott, Jacob G, and Vladimirsky, Alexander. “Optimizing adaptive cancer therapy: dynamic programming and evolutionary game theory”. In: Proceedings of the Royal Society B 287.1925 (2020), p. 20192454. doi: 10.1098/rspb.2019.2454.

[26] Henke, Erik, Nandigama, Rajender, and Ergün, Süleyman. “Extracellular Matrix in the Tumor Microenvironment and Its Impact on Cancer Therapy”. In: Frontiers in Molecular Biosciences 6 (Jan. 2020), p. 470149. issn: 2296889X. doi: 10.3389/FMOLB.2019.00160/BIBTEX.

[27] West, Jeffrey, et al. “Towards multidrug adaptive therapy”. In: Cancer research 80.7 (2020), pp. 1578–1589.

[28] Dillekås, Hanna, Rogers, Michael S., and Straume, Oddbjørn. “Are 90% of deaths from cancer caused by metastases?” In: Cancer Medicine 8.12 (Sept. 2019), p. 5574. issn: 20457634. doi: 10.1002/CAM4.2474.

[29] Gatenby, Robert A., Zhang, Jingsong, and Brown, Joel S. “First strike – second strike strategies in metastatic cancer: Lessons from the evolutionary dynamics of extinction”. In: Cancer research 79.13 (July 2019), p. 3174. issn: 15387445. doi: 10.1158/0008-5472.CAN-19-0807.

[30] Karagiannis, George S., Condeelis, John S., and Oktay, Maja H. “Chemotherapy-Induced Metastasis: Molecular Mechanisms, Clinical Manifestations, Therapeutic Interventions”. In: Cancer research 79.18 (Sept. 2019), p. 4567. issn: 15387445. doi: 10.1158/0008-5472.CAN-19-1147.

[31] Rosel, Daniel et al. “Migrastatics: Redirecting R&D in Solid Cancer Towards Metastasis?” In: Trends in cancer 5.12 (Dec. 2019), pp. 755–756. issn: 2405-8025. doi: 10.1016/J.TRECAN.2019.10.011. url: https://pubmed.ncbi.nlm.nih.gov/31813449/.

[32] Staňková, Kateřina. “Resistance games”. In: Nature Ecology & Evolution 2019 3:3 3.3 (Feb. 2019), pp. 336–337. issn: 2397-334X. doi: 10.1038/s41559-018-0785-y.

[33] Behrenbruch, Corina et al. “Surgical stress response and promotion of metastasis in colorectal cancer: a complex and heterogeneous process”. In: Clinical and Experimental Metastasis 35.4 (Apr. 2018), pp. 333–345. issn: 15737276. doi: 10.1007/S10585-018-9873-2/FIGURES/1.

[34] Cunningham, Jessica J. et al. “Optimal control to develop therapeutic strategies for metastatic castrate resistant prostate cancer”. In: Journal of theoretical biology 459 (2018), pp. 67–78.

[35] Gatenby, Robert A. and Brown, Joel S. “The Evolution and Ecology of Resistance in Cancer Therapy”. In: Cold Spring Harbor Perspectives in Medicine 8.3 (Mar. 2018). issn: 21571422. doi: 10.1101/CSHPERSPECT.A033415.

[36] Harmand, Nóemie, et al. “Evolution of bacteria specialization along an antibiotic dose gradient”. In: Evolution Letters 2.3 (June 2018), p. 221. issn: 20563744. doi: 10.1002/EVL3.52.

[37] Riihimäki, Matias, et al. “Clinical landscape of cancer metastases”. In: Cancer Medicine 7.11 (Nov. 2018), p. 5534. issn: 20457634. doi: 10.1002/CAM4.1697.

[38] Cree, Ian A. and Charlton, Peter. “Molecular chess? Hallmarks of anti-cancer drug resistance”. In: BMC Cancer 17.1 (Jan. 2017), pp. 1–8. issn: 14712407. doi: 10.1186/S12885-016-2999-1/FIGURES/1.

[39] Gandalovičová, Aneta, et al. “Migrastatics—Anti-metastatic and Anti-invasion Drugs: Promises and Challenges”. In: Trends in Cancer 3.6 (June 2017), pp. 391–406. issn: 24058033. doi: 10.1016/j.trecan.2017.04.008.

[40] Jones, Brandon C. et al. “Dual Targeting of Mesenchymal and Amoeboid Motility Hinders Metastatic Behavior”. In: Molecular cancer research : MCR 15.6 (June 2017), pp. 670–682. issn: 1557-3125. doi: 10.1158/1541-7786.MCR-16-0411. url: https://pubmed.ncbi.nlm.nih.gov/28235899/.

[41] Lambert, Arthur W., Pattabiraman, Diwakar R., and Weinberg, Robert A. “EMERGING BIOLOGICAL PRINCIPLES OF METASTASIS”. In: Cell 168.4 (Feb. 2017), p. 670. issn: 10974172. doi: 10.1016/J.CELL.2016.11.037.

[42] You, Li et al. “Spatial vs. non-spatial eco-evolutionary dynamics in a tumor growth model”. In: Journal of Theoretical Biology 435 (Dec. 2017), pp. 78–97. issn: 00225193. doi: 10.1016/j.jtbi.2017.08.022.

[43] Zhang, Jingsong et al. “Integrating evolutionary dynamics into treatment of metastatic castrate-resistant prostate cancer”. In: Nature Communications 8.1 (Nov. 2017), pp. 1–9. issn: 2041-1723. doi: 10.1038/S41467-017-01968-5.

[44] Jobe, Njainday Pulo et al. “Simultaneous blocking of IL-6 and IL-8 is sufficient to fully inhibit CAF-induced human melanoma cell invasiveness”. In: Histochemistry and cell biology 146.2 (Aug. 2016), pp. 205–217. issn: 1432-119X. doi: 10.1007/S00418-016-1433-8. url: https://pubmed.ncbi.nlm.nih.gov/27102177/.

[45] Obenauf, Anna C. and Massagué, Joan. Surviving at a Distance: Organ-Specific Metastasis. Sept. 2015. doi: 10.1016/j.trecan.2015.07.009.

[46] Wan, Liling, Pantel, Klaus, and Kang, Yibin. “Tumor metastasis: moving new biological insights into the clinic”. In: Nature medicine 19.11 (Nov. 2013), pp. 1450–1464. issn: 1546-170X. doi: 10.1038/NM.3391.

[47] Wu, Amy et al. “Cell motility and drug gradients in the emergence of resistance to chemotherapy”. In: Proceedings of the National Academy of Sciences of the United States of America 110.40 (Oct. 2013), pp. 16103–16108. issn: 00278424. doi: 10.1073/PNAS.1314385110/-/DCSUPPLEMENTAL.

[48] Sleeman, Jonathan and Steeg, Patricia S. “Cancer metastasis as a therapeutic target”. In: European Journal of Cancer 46.7 (May 2010), pp. 1177–1180. issn: 09598049. doi: 10.1016/j.ejca.2010.02.039.

[49] Gatenby, Robert A. “A change of strategy in the war on cancer”. In: Nature 459.7246 (2009), p. 508.

[50] Gatenby, Robert A. et al. “Adaptive Therapy”. In: Cancer research 69.11 (June 2009), p. 4894. issn: 00085472. doi: 10.1158/0008-5472.CAN-08-3658.

[51] Lage, H. “An overview of cancer multidrug resistance: a still unsolved problem”. In: Cellular and molecular life sciences 65.20 (Oct. 2008), pp. 3145–3167. issn: 1420-682X. doi: 10.1007/S00018-008-8111-5.

[52] Ratain, Mark J. and Mick, Robert. “Model-Guided Determination of Maximum Tolerated Dose in Phase I Clinical Trials: A Paradigm for Dose Selection in the Era of Targeted Therapies”. In: Journal of the National Cancer Institute 85.3 (1993), pp. 217–223. doi: 10.1093/jnci/85.3.217. url: https://academic.oup.com/jnci/article/85/3/217/983887.

